# Protein Kinase C Delta Regulates Mononuclear Phagocytes and Hinders Response to Immunotherapy in Cancer

**DOI:** 10.1101/2022.03.31.486620

**Authors:** Mehdi Chaib, Jeremiah Holt, Laura M. Sipe, Margaret S. Bohm, Sydney J. Clarice, Johnathan R. Yarbro, Ubaid Tanveer, T.J. Hollingsworth, QingQing Wei, Paul G. Thomas, D. Neil Hayes, Liza Makowski

## Abstract

Checkpoint immunotherapy unleashes T cell antitumor potential which has revolutionized cancer treatment showing unprecedented long-term responses. However, most patients do not respond to immunotherapy which often correlates with a dysfunctional or immunosuppressive myeloid compartment. The mononuclear phagocyte system (MPS) is a sub-class of myeloid cells comprising monocytes, macrophages and dendritic cells which plays a crucial role in tissue homeostasis. However, accumulating evidence suggests that mononuclear phagocytes contribute to all phases of tumorigenesis including orchestrating inflammatory events during de novo carcinogenesis, contribution to the progression of established tumors and promotion of resistance to checkpoint blockade. Thus, targeting the MPS could be an effective strategy to enhance checkpoint blockade efficacy and promote control of tumors. Here, we found that protein kinase C delta (PKCδ), a serine/threonine kinase, is abundantly expressed by mononuclear phagocytes in several human and mouse tumors. PKCδ^−/−^ mice were more resistant to growth of various cancers compared to wild-type mice and were more responsive to anti-PD-1 immunotherapy. Furthermore, we found that tumors from PKCδ^−/−^ mice harbor a Th-1-skewed immune landscape including increased antigen cross-presentation and T cell activation. Depletion of mononuclear phagocytes *in vivo* altered tumor growth in wild-type mice, but not in PKCδ^−/−^ mice. In addition, coinjection of PKCδ^−/−^-deficient M2-like macrophages with cancer cells into wild-type mice markedly delayed tumor growth and significantly increased intratumoral T cell activation compared to wild-type M2-like macrophages coinjected with cancer cells. Finally, intrinsic loss of PKCδ^−/−^ functionally reprogrammed macrophages and dendritic cells by promoting their antigen presenting and cross-presenting capacity and triggered type I and type II interferon signaling. Thus, PKCδ might be targeted to reprogram mononuclear phagocytes and augment checkpoint blockade efficacy.

## Introduction

Tumors develop in the context of a highly complex microenvironment that can greatly influence disease progression and response to therapy^1^. Immune cells are now widely recognized as a crucial component of the tumor microenvironment (TME) and are prognostic for clinical outcome in cancer patients^2^. Much of the field’s focus has been on approaches that reinvigorate adaptive immunity such as the use of immune checkpoint blockade (ICB) showing unprecedented durable responses^3^. Unfortunately, most patients do not respond to ICB for reasons that are still unclear^4,5^. One of the most important factors that contribute to immunotherapy resistance is the immunosuppressive nature of the TME which is largely shaped by innate immune cells, mainly myeloid cells^6^. This emphasizes the need to understand the signals that regulate myeloid cells in the TME.

Mononuclear phagocytes (MPs) comprising monocytes, macrophages, and dendritic cells (DCs) are a heterogenous innate immune cell population that plays a crucial role in host defense and tissue homeostasis^7^. However, MPs contribute to all phases of tumorigenesis including orchestrating inflammatory events during *de novo* carcinogenesis, contribution to the progression of established tumors, and promotion of resistance to ICB^8,9^. Due to their highly plastic nature, MPs often play opposing roles where they orchestrate antitumor responses on the one hand and promote immune suppression on the other^10^. Therefore, understanding the signals that regulate MP functional states may yield powerful targets to harness the anti-tumor potential of innate immunity to improve cancer immunotherapy response.

Monocytes are composed of two main subsets in mice and humans: classical and non-classical monocytes, and these cells are found predominantly in the circulation, bone marrow, and spleen^11^. Both monocyte subsets have been reported to have pro and antitumor properties^12–14^. Immature myeloid cells (iMCs), also defined as monocytic myeloid-derived suppressor cells (M-MDSCs) are another subset of the monocytic lineage and are highly immunosuppressive in cancer^15,16^. Monocytes and M-MDSCs express high levels of Ly6C in mice. Both cell types also play a role in tumor progression by differentiating into monocyte-derived macrophages or monocyte-derived DCs in the TME^10,17^. Tumor-associated macrophages represent the major tumor-infiltrating immune cell type in most solid tumors and are assumed to be tumor promoting^18^. DCs, on the other hand, are generally considered to be favorable for the antitumor response because of their remarkable antigen-presenting capacity^19–22^. However, DCs also have regulatory functions that limit antitumor immunity^23^. Consequently, identifying novel targets that can reprogram MPs in cancer are needed.

Protein kinase C delta (PKCδ), a serine/threonine kinase, is involved in several cellular processes including differentiation, apoptosis, cell survival and proliferation^24,25^. Autosomal recessive PKCδ deficiency in humans or genetic deletion of PKCδ in mice resulted in severe systemic autoimmunity^25,26,27^. In myeloid cells, loss of PKCδ resulted in impaired extracellular trap formation in neutrophils^28^ and decreased macrophage phagosomal clearance of microbes*^29,30^*. Whether PKCδ inhibits or promotes cancer cell growth is not clear from the litterature^31^. However, the role of PKCδ in antitumor immunity is largely unknown.

In this study, *Prkcd*^*−/−*^ mice displayed delayed tumor growth compared to wildtype (*Prkcd*^*+/+*^) mice using breast, lung, and melanoma cancer models. Delay of tumor growth was more significant in E0771 (breast) and LLC (lung) models which correlated with higher content of MPs in these tumors. The effects of PKCδ deficiency on tumor growth were associated with increased antigen-presentation and intratumoral CD8^+^ T cells, which expressed higher levels of activation markers protein death receptor 1 (PD-1), interferon gamma (IFNγ) and tumor necrosis factor alpha (TNFα).Overall, PKCδ deficiency induced a Th1-skewed immune response in the tumors. We also found PKCδ to be abundantly expressed by MPs across several human tumors using single cell RNAseq analysis of several publicly available databases. Importantly, depletion of MPs or MP-tumor cell co-injection experiments revealed that the effects of PKCδ deficiency on tumor growth and immune suppression were dependent on MPs. Mechanistically, intrinsic loss of PKCδ in MPs activated type I and II interferon signaling and enhanced their antigen-presenting and cross-presenting capability. Last, anti-PD-1 immunotherapy was more effective in PKCδ-deficient compared to wildtype tumor-bearing mice as evidenced by a marked delay in tumor growth and a significantly longer overall survival. In sum, PKCδ represents an attractive novel target to reprogram MPs and enhance ICB efficacy in cancer.

## Results

### PKCδ promotes tumor growth and immune suppression in mice

To explore the role of PKCδ in tumorigenesis, we implanted breast (E0771), lung (LLC) and melanoma (B16F10) syngeneic murine cancer cell lines into *Prkcd*^*+/+*^ and *Prkcd*^*−/−*^ mice. Compared to *Prkcd*^*+/+*^ mice, *Prkcd*^*−/−*^ mice exhibited a significant delay in tumor growth in E0771 and LLC models (Fig. 1A,B), but this effect was not significant in the B16F10 model (Fig. 1C). When we analyzed the intratumoral immune cell content of these tumors using flow cytometry, we found that E0771 and LLC tumors were abundantly infiltrated by MPs (27% and 35% of all viable cells, respectively), but not B16F10 tumors (1%) (Fig. 1D) which was consistent with previously published findings^15,32^. Decreased effect in the B16F10 model harboring fewer MPs suggests that PKCδ may primarily regulate MPs in cancer.

**Figure 1:**
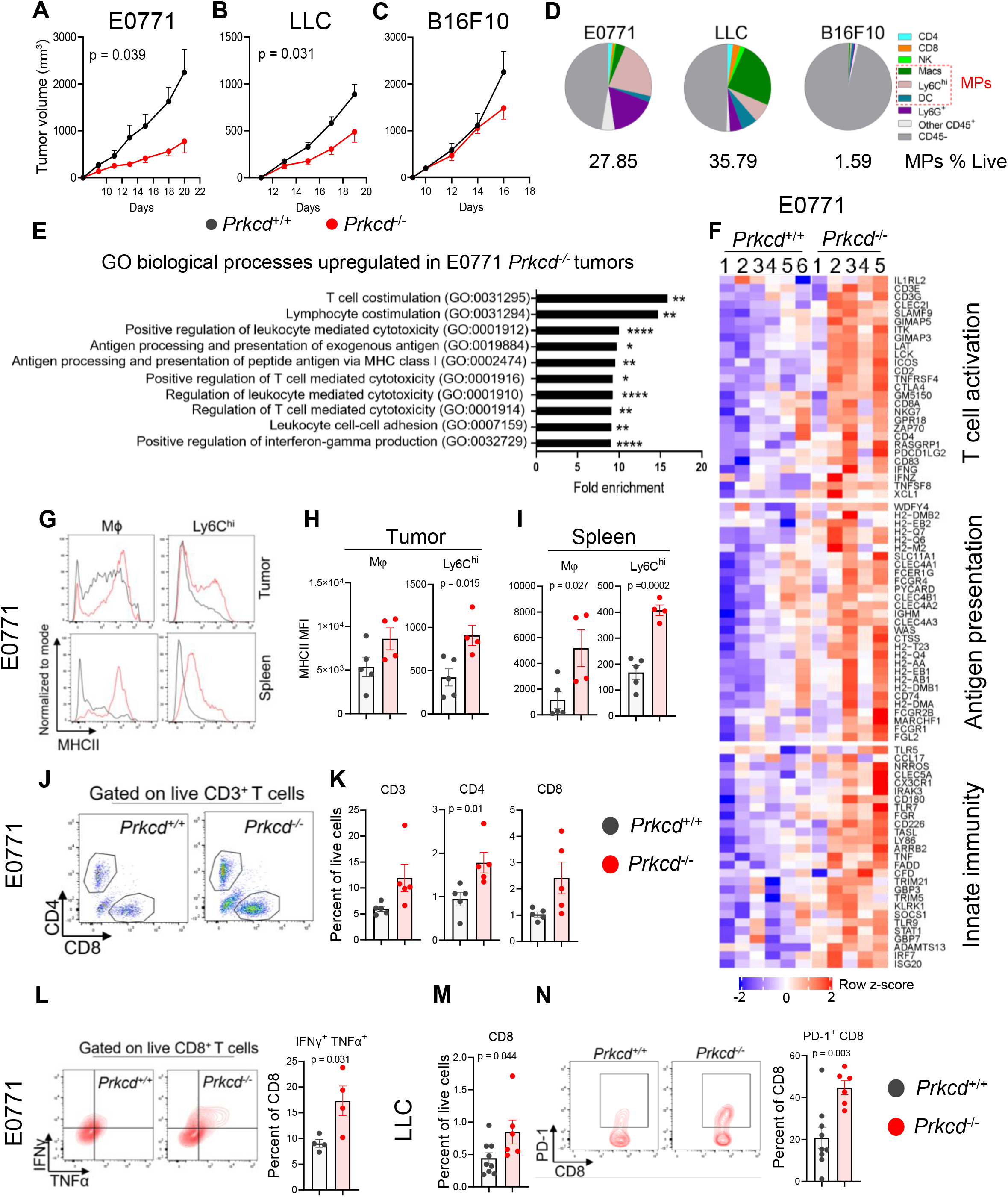
PKCδ promotes tumor growth and immune suppression. (A-C) Tumor volumes in *Prkcd*^+/+^ and *Prkcd*^−/−^ mice orthotopically injected with **(A)** E0771 breast cancer cells (n = 5-6 biological replicates) or subcutaneously injected with **(B)** LLC lung cancer cells (n = 7-9 biological replicates) and **(C)** B16F10 melanoma cancer cells (n = 5 biological replicates). Two-way ANOVA was used. **(D)** Immune and non-immune cell composition of E0771, LLC, and B16F10 tumors and proportions of mononuclear phagocytes (MPs) as analyzed by flow cytometry (n = 4 biological replicates). **(E)** Gene Ontology (GO) analysis of the genes that were uniquely upregulated in E0771 *Prkcd*^−/−^ tumors. Bonferroni correction for multiple testing was used (*P<0.05, **P<0.01, ****P<0.0001). **(F)** Heatmap of median-centered mRNA expression of genes involved in the immune response in tumors from *Prkcd*^+/+^ and *Prkcd*^−/−^ mice (n = 5-6 biological replicates). **(G-I)** MHCII expression in macrophages and Ly6C^hi^ cells from **(H)** E0771 tumors and **(I)** spleens from same tumor bearing mice as quantified by mean fluorescence intensity (MFI, n = 4-5 biological replicates). (**J-K**) Flow cytometry analysis of CD3^+^, CD4^+^, and CD8^+^ T cell content in E0771 tumors reported as frequency of live cells (n = 5 biological replicates). **(L)** Frequency of IFNγ^+^ TNFα^+^ CD8^+^ T cells in E0771 tumors (n = 4 biological replicates). **(M)** CD8^+^ T cell and **(N)**PD-1^+^ CD8^+^ T cell content in LLC tumors (n = 6-9 biological replicates). Unpaired Student’s T-test was used in flow cytometry analysis (p < 0.05 was considered significant). Data are shown as mean ± SEM.

We next examined the effect of PKCδ deficiency on gene regulation by bulk RNA sequencing (RNAseq) and analysis of differentially expressed genes (DEGs) in E0771 tumors. There were 473 significantly upregulated and 240 significantly downregulated genes in *Prkcd*^*−/−*^ versus *Prkcd*^*+/+*^ tumors (Supplementary Fig. 2A). Gene ontology (GO) analysis of genes upregulated in *Prkcd*^*−/−*^ revealed enhanced immunostimulatory responses (such as T cell activation, interferon gamma (IFNγ) signaling and antigen presentation) (Fig. 1E). In addition, genes involved in antigen presentation, innate immunity and T cell activation were elevated in *Prkcd*^*−/−*^ tumors compared to *Prkcd*^*+/+*^ tumors (Fig. 1F). Similarly, gene set enrichment analysis (GSEA) revealed significant enrichment for multiple immune-related GO pathways in *Prkcd*^*−/−*^ tumors including T cell activation, antigen processing and presentation, innate immune response, and inflammatory response (Supplementary Fig. 2B-E).

Flow cytometry analysis revealed enhanced expression of major histocompatibility complex class II (MHCII) in macrophages and monocytes/iMCs (Ly6C^hi^ cells) from tumors (Fig. 1G,H) and spleens (Fig. 1G,I) of *Prkcd*^*−/−*^ compared to *Prkcd*^*+/+*^ E0771 tumor-bearing mice which is suggestive of enhanced maturation and antigen-presenting capacity of these cells. We also observed a significant increase in T cell content (total CD3^+^ T cells, CD4^+^ and CD8^+^ T cells) in E0771 *Prkcd*^*−/−*^ tumors (Fig. 1J,K) compared to *Prkcd*^*+/+*^ tumors. Importantly, there was a significant increase in CD8^+^ T cell activation (IFNγ^+^ TNFα^+^) in E0771 *Prkcd*^*−/−*^ tumors (Fig. 1L). In *Prkcd*^*−/−*^ LLC tumors, CD8^+^ T cell content (Fig. 1M) and PD-1^+^ CD8^+^ T cells (Fig. 1N) were significantly elevated compared to *Prkcd*^*+/+*^ tumors. Cumulatively, these results demonstrate that PKCδ deficiency restricts the growth of tumors that are highly infiltrated by MPs which suggested that this restraint may be associated with changes in infiltrating MPs that may impact T cell responses.

### PKCδ is abundantly expressed by mononuclear phagocytes

Since an immune response was required for tumor regression in *Prkcd*^*−/−*^ mice, we asked which cells express high levels of PKCδ in tumors but also in organs at steady state. First, we investigated PKCδ expression at a cellular level in several human tumors using single-cell RNAseq analysis of publicly available datasets. We found that a substantial fraction of MPs abundantly expressed PRKCD (PKCδ gene) in human triple negative breast cancer (TNBC)^33^ (Fig. 2A,B), melanoma^34^ (Fig. 2C,D), renal cell carcinoma^35^, colon cancer^36^, and glioblastoma^37^ tumors (Supplementary Fig. 3A-C, respectively). PKCδ was also abundantly expressed by MPs at steady state in human peripheral blood mononuclear cells (PBMC, Broad/Boston and MtSinai/NYC) (Supplementary Fig. 3D) and mouse CD45^+^ splenocytes (Immgen labs, Supplementary Fig. 3E).

**Figure 2:**
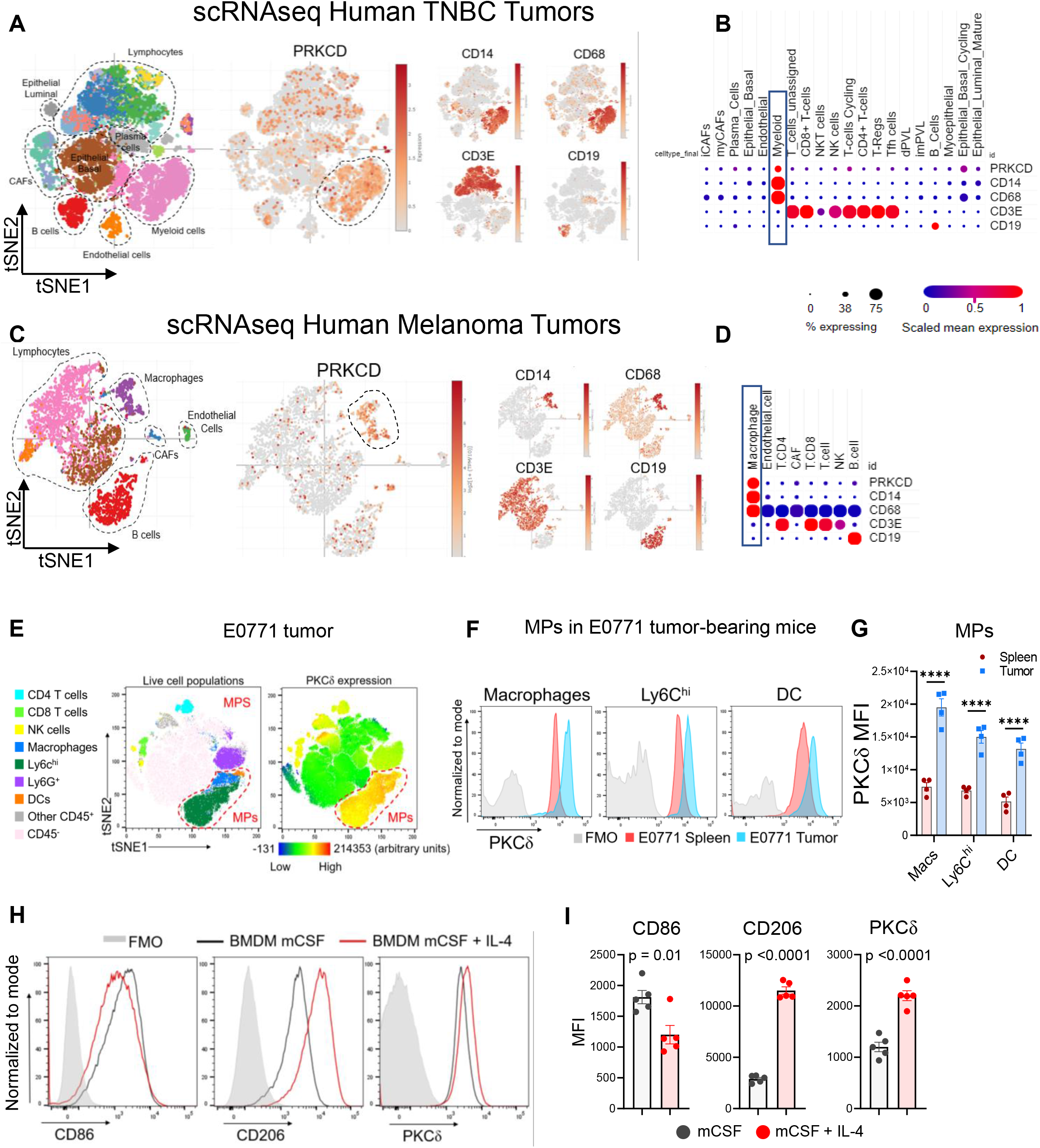
PKCδ is abundantly expressed by mononuclear phagocytes in cancer. (**A-D**) tSNE plots of single cell RNAseq showing major cell types, PRKCD mRNA expression, expression of monocyte/macrophage markers CD14 and CD68, T cell marker CD3E, and B cell marker CD19 in human **(A)** TNBC tumors (Wu et al., EMBO J 2020)^33^ and **(C)** melanoma tumors (Jerby-Arnon et al., Cell 2018)^34^. Percent of cells expressing the gene of interest and scaled mean expression is quantified with MPs highlighted in blue box **(B&D)**. **(E)** Representative tSNE dimensionality reduction plot showing concatenated flow cytometry analysis of live cell populations in E0771 tumors and PKCδ expression. Mononuclear phagocytes (MPs) are highlighted (n = 4 biological replicates). **(F)** Representative histograms and **(G)** MFI quantification of PKCδ expression in spleen and tumor MPs of E0771 tumor-bearing mice as quantified by MFI (n = 4 biological replicates). Paired T-test was used (****P<0.0001). **(H-I)** Wildtype BMDMs were polarized with 20 ng/ml of mouse recombinant IL-4 for 24h (red line) or left untreated as vehicle control (black line), with FMO control (grey). **(H)** Representative histogram and **(I)** MFI quantification of M1 marker CD86, M2 marker CD206 and PKCδ (n = 5 biological replicates). Unpaired Student’s T-test (p < 0.05 was considered significant). Data are shown as mean ± SEM.

Next, we checked PKCδ protein expression in spleen and tumor cell populations from E0771 tumor-bearing mice using flow cytometry. We found that PKCδ was predominantly expressed by MPs in E0771 tumors (Fig. 2E). In addition, MPs from E0771 tumor-bearing mice had significantly higher expression of PKCδ in the tumors compared to the spleen (Fig. 2F,G). Of note, myeloid cells from tumors are more immunosuppressive than their counterparts in the spleens^38^. Thus, PKCδ expression correlated with more immunosuppressive MPs which hints to a potential role in promoting immune suppression in MPs. Next, we checked PKCδ expression in M2-like (alternatively activated) polarized bone marrow-derived macrophages (BMDMs) which are known to be immunosuppressive and tumor promoting^18^. M2-like BMDMs expressed lower levels of the M1 marker CD86 and higher levels of the M2 marker CD206, as expected. Interestingly, we also found that PKCδ expression was significantly higher in M2-like BMDMs compared to non-polarized BMDMs (Fig. 2H,I). Taken together, these findings suggest that PKCδ may be a critical controller of MP regulatory or immunosuppressive states.

### PKCδ deficiency impairs tumor growth and immune suppression via mononuclear phagocytes

To investigate whether PKCδ deficiency in MPs is required for tumor repression and T cell activation, we first depleted MPs in *Prkcd*^*+/+*^ and *Prkcd*^*−/−*^ E0771 tumor-bearing mice using a combination of anti-Ly6C monoclonal antibody and clodronate liposomes^39^ (Fig. 3A). In accordance with previous reports^40–42^, we observed that MP depletion significantly delayed tumor growth in wildtype mice (*Prkcd*^*+/+*^) (Fig. 3B,D). By contrast, MP depletion in *Prkcd*^*−/−*^ mice did not delay tumor growth, but instead promoted tumor growth to an extent that is comparable with *Prkcd*^*+/+*^ mice (Fig. 3C,D). These results indicate that PKCδ deficiency likely reprograms MPs from a protumor phenotype to an anti-tumor phenotype.

**Figure 3:**
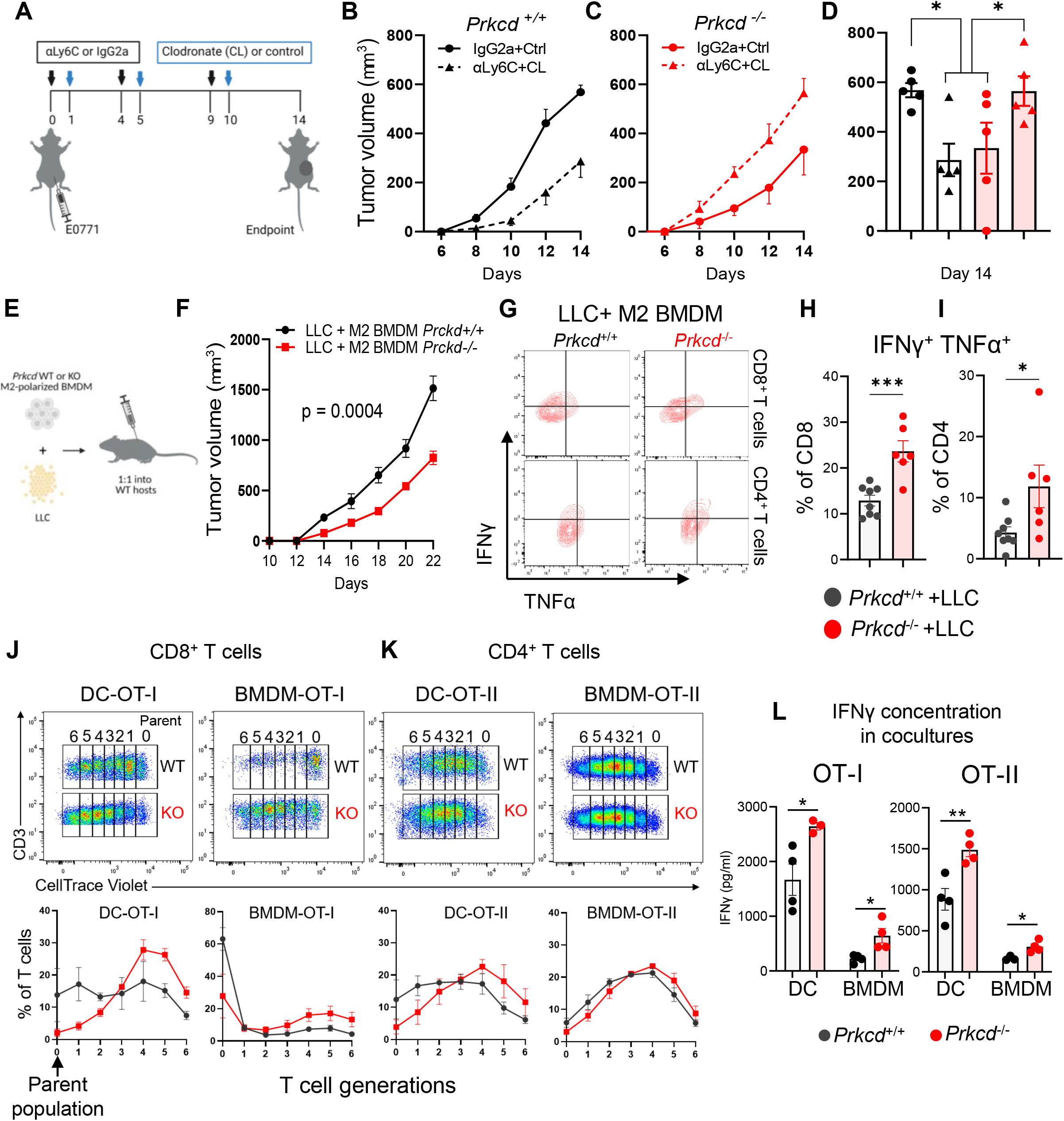
Loss of PKCδ impairs tumor growth and immune suppression *via* mononuclear phagocytes. *Prkcd*^+/+^ and *Prkcd*^−/−^ mice bearing E0771 tumors were treated with anti-Ly6C or IgG2a mAb (100μg/mouse) followed by clodronate or control liposome (200μl/mouse) as shown in experimental outline **(A)**. Tumor volume in **(B)** *Prkcd*^+/+^ and **(C)** *Prkcd*^−/−^ mice and **(D)** tumor volumes at day 14 in *Prkcd*^+/+^ and *Prkcd*^−/−^ mice treated as in **(A)** are shown. One-way ANOVA with multiple comparisons with Tukey correction was used (*P<0.05). **(E-I)** LLC cells were co-injected with M2-polarized BMDMs (20ng/mL of IL-4 for 24h) at a 1:1 ratio into wildtype mice. **(E)** Experimental outline and **(F)** tumor volume (n = 8 biological replicates). Two-way ANOVA was used. **(G-I)** Frequencies of IFNγ^+^ TNFα^+^ **(H)** CD8^+^ and **(I)** CD4^+^T cells in tumors from **(E-F)** was quantified and compared using unpaired Student’s T-test. **(J-L)** *Prkcd*^+/+^ or *Prkcd*^−/−^ bone marrow DCs and BMDMs were incubated with OVA (10μg/mL) overnight before coculture with CellTrace Violet-labelled CD8^+^ and CD4^+^ T cells isolated from OT-I and OT-II mice, respectively, for 3 days at a 2:1 T cell-DC/BMDM ratio. The individual peaks of CellTrace Violet dilution are highlighted as T cell generations ranging from 0 (parent population) to 6 (last daughter generation) and graphical representation of fractions of T cells in each peak in gated **(J)** CD8^+^ T cells and **(K)** CD4^+^ T cells. **(L)** IFNγ concentration in OT-I and OT-II co-culture supernatants from (J-K) was determined by ELISA (n = 3-4 biological replicates). Unpaired Student’s T-test (*P<0.05, **P<0.01). Data are shown as mean ± SEM.

We next investigated whether PKCδ deficiency in M2-like BMDMs decreases their tumor-promoting and T cell suppressive activity^43,44^ (Fig. 3E). Cancer cells (LLC) coinjected with *Prkcd*^*−/−*^M2-like BMDMs had a significant delay in tumor growth compared to LLC coinjected with *Prkcd*^*+/+*^ M2-like BMDMs (Fig. 3F). Importantly, we observed a significant increase in the activation (IFNγ^+^ TNFα^+^) of CD8^+^ (Fig. 3G,H) and CD4^+^ (Fig. 3G,I) T cells from *Prkcd*^*−/−*^ M2-like BMDMs + LLC tumors compared to *Prkcd*^*+/+*^ M2-like BMDMs + LLC tumors. Collectively, our findings suggest that PKCδ plays a critical role in controlling MP-induced effector T cell suppression and subsequent tumor promotion.

### PKCδ deficiency enhances antigen-presenting and cross-presenting capacity in mononuclear phagocytes

Antigen presentation to CD4^+^ T cells and antigen cross-presentation to CD8^+^ T cells are hallmark properties of antigen-presenting cells (APCs) to mount an effective antitumor immune response^45^. We pulsed BMDMs and DCs isolated from *Prkcd*^*+/+*^ and *Prkcd*^*−/−*^ mice with ovalbumin (OVA) before incubation with H-2K^b^-OVA peptide-specific T cell receptor (TCR) transgenic OT-I CD8^+^ T cells or OT-II CD4^+^ T cells, and measured T cell proliferation by analyzing the dilution of CellTrace Violet proliferation dye. We found that *Prkcd*^*−/−*^ BMDMs and DCs were superior at inducing OT-I CD8^+^ (Fig. 3J) and OT-II CD4^+^ (Fig. 3K) T cell proliferation compared to *Prkcd*^*+/+*^ BMDMs and DCs. In addition, PKCδ deficiency in BMDMs and DCs significantly increased IFNγ production in both OT-I and OT-II coculture supernatants as evaluated by ELISA (Fig. 3L). Findings herein indicate that PKCδ is critical in regulating MP-mediated T cell activation.

### Intrinsic loss of PKCδ triggers type I and type II IFN signaling in mononuclear phagocytes

To understand how PKCδ regulates MPs, we performed transcriptome analysis using RNAseq data from *Prkcd*^*−/−*^ and *Prkcd*^*+/+*^ M1-like BMDMs, stimulated DCs (DCstim), iMCs (Supplementary Fig. 4A,B), and whole E0771 tumors. We identified 552, 754, and 219 genes that were upregulated in *Prkcd*^*−/−*^ M1 BMDMs, DCstim, and iMCs, respectively, whereas 391, 905, and 186 genes were downregulated in these cells, respectively (Fig. 4A,E,I). Gene set enrichment analysis (GSEA) revealed that hallmark pathways that are associated with a proinflammatory phenotype such as response to IFNα/γ and inflammatory response were significantly enriched in M1 BMDMs and DCstim compared to unstimulated BMDM and DCs, respectively, suggesting that these cells have been successfully polarized towards a proinflammatory phenotype (Supplementary Fig. 4C,D). Interestingly, GSEA revealed that response to IFNα/γ hallmark pathways were consistently highly enriched in *Prkcd*^*−/−*^ M1 BMDMs (Fig. 4B–D), DCstim (Fig. 4F–H), iMCs (Fig. 4J–L) and E0771 tumors (Fig. 4M–O) suggesting that Type I and type II interferon signaling pathways are triggered in PKCδ-deficient MPs. We also observed elevated expression of genes involved in the response to type I interferon in *Prkcd*^*−/−*^ E0771 tumors compared to *Prkcd*^*+/+*^ tumors (Supplementary Fig. 5). By contrast, pathways significantly enriched in *Prkcd*^*+/+*^ M1 BMDM, DCstim, iMCS, and E0771 tumors included hallmark gene sets involved in promotion of tumor growth and metastasis such as epithelial to mesenchymal transition (EMT) and angiogenesis as well as anti-inflammatory pathways such as bile acid metabolism and coagulation^46–48^, which were consistently highly enriched in *Prkcd*^*+/+*^ M1 BMDM, DCstim, iMCS, and E0771 tumors (Supplementary Fig. 6A-E).

**Figure 4:**
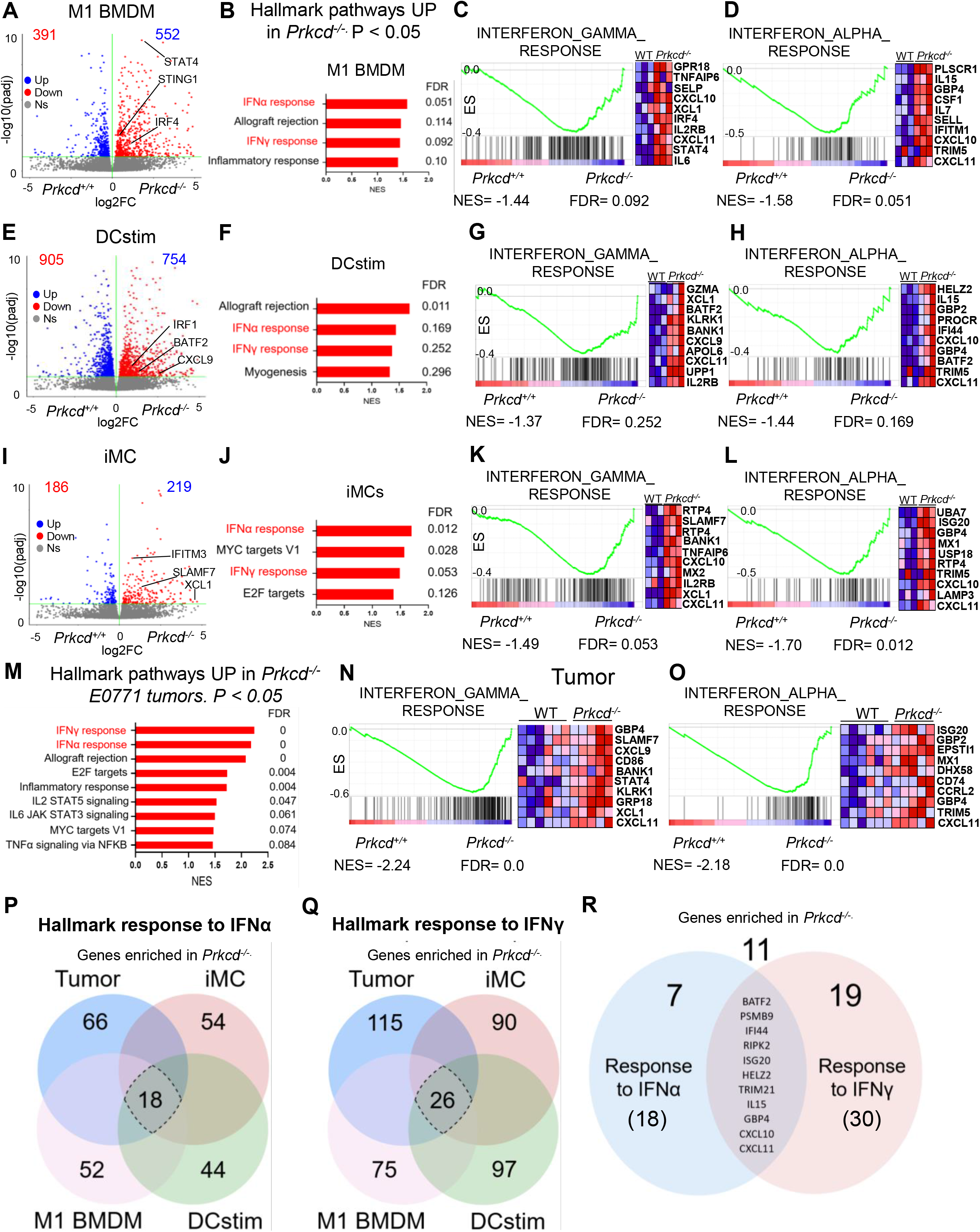
Intrinsic loss of PKCδ triggers type I and type II IFN signaling in mononuclear phagocytes. (**A-D)** M1 BMDMs were obtained by stimulating BMDMs with IFNγ (20ng/mL) and LPS (100ng/mL) for 24h. **(A)** Volcano plot for all differentially expressed genes between *Prkcd*^+/+^ and *Prkcd*^−/−^ M1 BMDMs is shown. **(B)** Gene set enrichment analysis (GSEA) of hallmark gene sets (H.all) from the Molecular Signatures Database of the Broad Institute is reported, showing the most significantly enriched gene sets in *Prkcd*^−/−^ M1 BMDMs and their normalized enrichment score (NES). GSEA plots of the **(C)** interferon gamma response and **(D)** interferon alpha response in *Prkcd*^+/+^ and *Prkcd*^−/−^ M1 BMDMs are shown. The top 10 enriched genes by enrichment score in *Prkcd*^−/−^ relative to *Prkcd*^+/+^ from each category are shown. **(E-H)** Stimulated DCs (DCstim) were obtained by stimulating bone marrow DCs with LPS (100ng/mL) and agonistic CD40 mAb (5μg/mL) for 24h. Control DCs were treated with IgG2a (5μg/mL). **(E)** Volcano plot for all differentially expressed genes between *Prkcd*^+/+^ and *Prkcd*^−/−^ DCstim is shown. **(F)** GSEA of hallmark gene sets showing the most significantly enriched gene sets in *Prkcd*^−/−^ DCstim is presented. GSEA plot of the **(G)** interferon gamma response and **(H)** interferon alpha response in *Prkcd*^+/+^ and *Prkcd*^−/−^ DCstim is shown. **(I-L)** Immature myeloid cells (iMC) were generated by culturing bone marrow cells with GM-CSF (40ng/mL) and IL-6 (40ng/mL) for 6 days. **(I)** Volcano plot for all differentially expressed genes between *Prkcd*^+/+^ and *Prkcd*^−/−^ iMCs is shown. **(J)** GSEA of hallmark gene sets showing the most significantly enriched gene sets in *Prkcd*^−/−^ iMCs. GSEA plot of the **(K)** interferon gamma response and **(L)** interferon alpha response in *Prkcd*^+/+^ and *Prkcd*^−/−^ iMCs. **(M)** GSEA of hallmark gene sets showing the most significantly enriched gene sets in *Prkcd*^−/−^ E0771 tumors compared to *Prkcd*^+/+^. GSEA plot of the **(N)**interferon gamma and **(O)** interferon alpha response in *Prkcd*^+/+^ and *Prkcd*^−/−^ E0771 tumors. **(P-R)** Venn diagrams of enriched genes in *Prkcd*^−/−^ tumors, M1 BMDMs, DCstim and iMCs for the hallmark gene sets **(P)** response to interferon alpha and **(Q)** response to interferon gamma relative to *Prkcd*^+/+^ controls. **(R)** Venn diagrams of commonly enriched genes between *Prkcd*^−/−^ tumors, M1 BMDMs, DCstim, and iMCs for response to interferon alpha (18 common genes) and response to interferon gamma (26 common genes). (n = 3 biological replicates for all MPs and n = 5-6 for E0771 tumors). For GSEA hallmark gene sets, nominal p value was less than 0.05 for all shown pathways. For volcano plots, differentially expressed genes with an adjusted p value less than 0.1 were considered.

The gene sets induced by Type I and Type II interferons overlap considerably, and both are essential to induce T cell activation and protective immunity^49^. We therefore investigated commonly enriched genes between *Prkcd*^*−/−*^ M1 BMDM, DCstim, iMCS, and E0771 tumors from both hallmark gene sets response to IFNα (18 genes) (Fig. 4P) and response to IFNγ (26 genes) (Fig. 4Q). We found 11 overlapping genes between the two interferon gene sets (Fig. 4R) which may represent the most commonly upregulated interferon responsive genes in PKCδ deficient MPs. Taken together, our findings reveal a potential role of PKCδ in promoting protumor and anti-inflammatory pathways while repressing type I and II interferon pathways in MPs.

### PKCδ deficiency enhances anti-PD-1 therapy

The antitumor effect observed in *Prkcd*^*−/−*^ mice prompted us to determine whether PKCδ deficiency can improve responsiveness to ICB. We chose the LLC tumor model previously reported as being relatively resistant to ICB^32,50^. LLC tumor-bearing *Prkcd*^*+/+*^ and *Prkcd*^*−/−*^ mice were treated with anti-PD-1 or IgG2a control as outlined in Fig. 5A. Although we observed a moderate but significant reduction in tumor growth in *Prkcd*^*+/+*^ mice treated with anti-PD-1 compared to IgG2a-treated *Prkcd*^*+/+*^ mice, combination of PKCδ deficiency and anti-PD-1 synergistically delayed tumor growth (Fig. 5B,C). Notably, *Prkcd*^*−/−*^ mice treated with anti-PD-1 had a significantly prolonged overall survival compared to other groups (Fig. 5D). Taken together, our findings indicate that PKCδ may represent a promising novel target to improve responsiveness to ICB.

**Figure 5:**
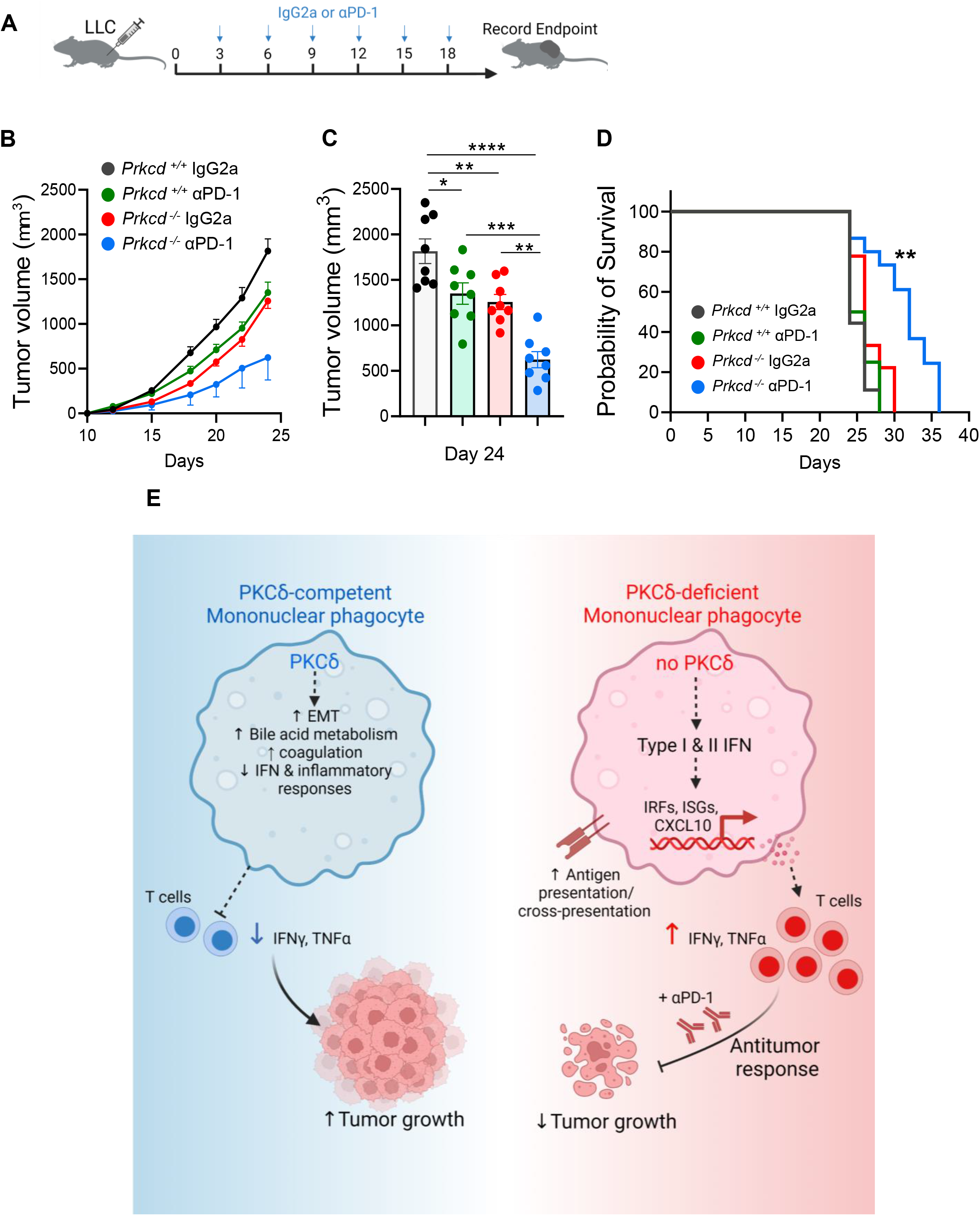
Loss of PKCδ improves response to anti-PD-1 therapy. LLC tumor-bearing *Prkcd*^+/+^ and *Prkcd*^−/−^ mice were treated with anti-PD-1 (200μg/mouse) or IgG2a (200μg/mouse) every 3 days as shown in **(A)**. **(B)** Tumor volume over time until day 24, **(C)** tumor volume at day 24 and **(D)** Kaplan-Meier survival curves of tumor-bearing mice are shown (n = 8 mice per group). **(F)** Proposed model of PKCδ function in mononuclear phagocytes and tumor progression. One-way ANOVA with multiple comparisons with Tukey’s correction was used to compare tumor volumes (*P<0.05, **P<0.01, ***P<0.001, ***P<0.0001). Data are shown as mean ± SEM. Log-rank (Mantel-Cox) test was used to determine statistical significance for survival of mice in E (**P<0.01, ***P<0.001, ***P<0.0001).

## Discussion

Resistance to ICB poses a major challenge to the therapeutic management of patients with solid tumors. Currently, most research efforts aiming at improving immunotherapy outcomes focus on T cells. However, given that innate immunity plays a critical role in orchestrating adaptive immunity, incorporating both arms of the immune system could be a more effective strategy to improve immunotherapy efficacy. In this study, we identified an immune evasion mechanism by which MPs are wired to suppress the antitumor immune response *via* PKCδ signaling. In this context, PKCδ acts as an innate immune checkpoint. We show that genetic deletion of PKCδ curbs tumor growth and promotes T cell tumor infiltration and activation in preclinical cancer models that have high MP content in their tumors. We also show that loss of PKCδ in MPs had a profound effect on the overall transcriptional program which resulted in their reprogramming to an antitumor phenotype. PKCδ-deficient MPs activate Type I and II interferon signaling which are often required for mounting an antitumor immune response^51^. These results highlight two key points: (i) the importance of MPs in controlling antitumor immunity and (ii) that PKCδ is a critical driver of MP phenotype in the TME and a potential novel target in cancer immunotherapy.

Although ICB has recently revolutionized cancer treatment, most patients fail to respond due to several factors, one of which is the establishment of a suppressive TME rich in myeloid cells^6^. Thus, efforts are currently ongoing to identify novel myeloid targets to complement ICB. Some of these approaches focus on blocking suppressive MP cell recruitment to the TME, inhibiting their pro-tumoral functions, or restoring their immunostimulatory properties. Among others, these approaches include inhibition of phosphoinositide 3-kinase gamma (PI3Kγ)^52^ and colony stimulating factor 1 receptor (CSF-1R)^53^, as well as blockade of TREM2^6^ and TAM receptors (Tyro3, Axl, and MerTK)^54^. In our study, we found that PKCδ deficiency combined with anti-PD-1 markedly delayed tumor growth and significantly extended the survival of LLC tumor-bearing mice. Thus, PKCδ inhibition provides a novel therapeutic approach that broadens the arsenal of myeloid cell targeting in tumors. Although several studies claim the existence of PKCδ specific inhibitors, one must be cautious using these inhibitors to specifically target the delta isoform of the PKC family due to several potential challenges^55^. One of these challenges is off-target effects such as inhibition of other PKC isoforms that share similarities with the PKCδ protein structure. Some of these PKC isozymes may play contrasting physiological roles to PKCδ which can result in dampening the desired effects of PKCδ inhibition^56^. Therefore, developing therapeutic tools to specifically inhibit PKCδ may represent a promising therapeutic strategy to enhance immunotherapy efficacy in cancer patients.

PKCδ is a serine/threonine kinase of the novel PKC sub-family and can be activated by stimulation with diacylglycerol leading to PKCδ phosphorylation and activation of downstream targets^24^. PKCδ is involved in a myriad of cellular processes involving apoptosis, proliferation, and cell survival in a variety of cell types including immune cells^24,25^. In the hematopoietic compartment, studies have shown that genetic deletion of PKCδ resulted in systemic autoimmunity which correlated with accumulation of autoreactive B cells in PKCδ knockout mice^26,27^. Similarly, patients with autosomal recessive PKCδ deficiency were severely autoimmune and suffered from systemic lupus erythematosus^25^. In myeloid cells, previous work demonstrated that loss of PKCδ resulted in a defective reactive oxygen species production and impaired extracellular trap formation in neutrophils^28^ and decreased macrophage phagosomal clearance of *Listeria monocytogenes* and *Mycobacterium tuberculosis^29,30^.* Although PKCδ is widely characterized as a pro-apoptotic protein in cancer cells, much of the literature is still conflicted as to whether PKCδ inhibits or promotes cancer cell growth^31^. Our work aligns with previous studies by demonstrating that PKCδ plays a crucial role in regulating the immune response by acting as a brake on MP activation. Although this effect may be desirable at steady state to prevent autoimmunity^57^, it is however detrimental in cancer where an immune response is necessary to control tumors. In our study, we show that PKCδ is consistently and abundantly expressed by MPs across several human tumors. We also found that PKCδ is variably expressed by B cells and cancer cells depending on the tumor or organ Type. Therefore, future studies are needed to decipher the role of PKCδ in other hematopoietic and non-hematopoietic cells in cancer.

The underlying molecular mechanisms by which PKCδ dampens MP activation remain unclear. Our data suggest that this effect may be achieved by PKCδ activation of downstream pathways such as coagulation, bile acid metabolism, and EMT all of which promote the protumor and/or anti-inflammatory phenotype in MPs. In our study, PKCδ was shown to repress Type I and II interferon pathways, which are essential in orchestrating an effective T cell-mediated antitumor immune response^58^. However, the exact molecular interactions by which PKCδ represses interferon signaling will be subject of future investigations. While the evidence provided herein is supportive towards targeting MP PKCδ, a limitation is that our models were syngeneic transplants. It remains to be seen whether targeting PKCδ will be a successful strategy against spontaneous tumorigenesis and more advanced disease. Another limitation is that constitutive PKCδ deficiency might trigger direct or indirect compensatory responses of MPs that impact tumor growth. It will be interesting to determine whether acute inhibition of PKCδ using specific pharmacological agents results in complete control of tumor progression. Future studies using specific PKCδ inhibitors will be important to demonstrate how pharmacological inhibition of PKCδ in MPs interferes with signaling pathways and how it impacts tumor growth. In conclusion, this report demonstrates that PKCδ is a key driver of MP protumor phenotype in the TME, revealing a key novel target for cancer immunotherapy.

## Materials and Methods

### Reagents

All reagents were obtained from Sigma-Aldrich (St. Louis, MO) unless otherwise noted. Fetal bovine serum (FBS, Gibco, Waltham, MA), 100X L-glutamine, 100X penicillin/streptomycin HyClone (Pittsburgh, PA), and Gibco 100X antibiotic mix were obtained from Thermo Fisher (Waltham, MA). RPMI 1640, DMEM and Matrigel are from Corning (Tewksbury, MA). Mouse recombinant GM-CSF, IL-6, IL-4, M-CSF and FLT3L were obtained from Biolegend (San Diego, CA). Ovalbumin was obtained from Thermo Fisher. Mouse IFN gamma ELISA kit was obtained from R&D Systems (Minneapolis, MN). Mouse CD4^+^ T cell isolation kit and CD8^+^ T cell isolation kit were obtained from Miltenyi Biotec (Auburn, CA). Clodronate and control liposomes were obtained from Liposoma (Amsterdam, The Netherlands)^59^. In vivo anti-mouse CD40, anti-mouse PD-1, anti-mouse Ly6C monoclonal antibodies and their controls (rat IgG2a) were all obtained from BioxCell (Lebanon, NH). Knockout-validated PKCδ antibody and PE/Cy7 conjugation kit were obtained from Abcam (Cambridge, UK). Flow cytometry antibodies, compensation beads, and reagents are described in Supplemental Table 1 ((Tonbo, (San Diego, CA), Thermo Fisher and Biolegend.

### Animals

Animal studies were performed with approval and in accordance with the guidelines of the Institutional Animal Care and Use Committee (IACUC) at the University of Tennessee Health Science Center and in accordance with the National Institutes of Health Guide for the Care and Use of Laboratory Animals. All animals were housed in a temperature-controlled facility with a 12-h light/dark cycle and ad libitium access to food and water. *Prkcd*^−/−^ mice were a kind gift from Dr. Zheng Dong at Augusta University, Augusta GA and were generated as previously described^60^. After genotyping, only age- and sex-matched wildtype *Prkcd*^*+/+*^ and *Prkcd*^−/−^ mice were used in experiments. C57BL/6J (Stock No: 000664) mice were purchased from Jackson Laboratories (Bar Harbor, ME). For OT-I CD8^+^ T cell and OT-II CD4^+^ T cell studies, spleens from transgenic mice expressing the MHC-I restricted T cell receptor specific for the octamer SIINFEKL peptide ovalbumin257-264 (OT-I mice) and MHC-II restricted T cell receptor for the octamer SIINFEKL peptide ovalbumin 323-339 (OT-II mice) were a kind gift from Dr. Hongbo Chi at St Jude Children’s Research Hospital, Memphis TN.

### Tumor mouse models

8–12 week-old sex-matched *Prkcd*^*+/+*^ or *Prkcd*^−/−^ mice were used in *in vivo* experiments. E0771-luciferase (luc), a kind gift from Dr. Hasan Korkaya, Augusta University, is a murine adenocarcinoma breast cancer cell line that was originally isolated from a C57BL/6 mouse spontaneous tumor. Cells were cultured and injected as we previously described^3^. Briefly, cells were cultured in RPMI containing 10% FBS, 100 UI/mL of penicillin, and (100 μg/ml) streptomycin in a humidified chamber at 37°C under 5% CO_2_. E0771 cells were implanted into the left fourth mammary fat pad of 8-week-old C57BL/6J females at 250,000 cells in 100μl of 25% Matrigel. Murine Lewis Cell Carcinoma (LLC) cells (10^6^ cells unless otherwise specified), a kind gift from Dr. James A. Carson from UTHSC, Memphis TN and murine B16F10 melanoma cells (3 × 10^5^ cells), a kind gift from Dr. Hongbo Chi at St. Jude Children’s and Research Hospital in Memphis TN were cultured in DMEM as above and were subcutaneously implanted in PBS into the right flank of male mice as noted. Tumor growth was monitored by measuring the length and width of the tumor using digital calipers. Tumor volume was calculated using the following formula^61^: Volume = (width)^2^ ×□(length)/2.

### Anti-PD-1 tumor studies

8–12-week-old female *Prkcd*^*+/+*^ or *Prkcd*^−/−^ mice were implanted with LLC cells (2 × 10^5^) as above. Mice from each genotype were randomized then treated with six doses of anti-PD-1 or rat IgG2a (200μg/mouse) every three days starting at day 3. Survival events were scored when tumor volume reached >2000 cm^3^, or when mice had moribund appearance, reached endpoint per ICUC guidelines, or per absolute survival.

### *In vivo* mononuclear phagocyte depletion studies

Mononuclear phagocyte depletion experiments were conducted as previously described^39^ with some modifications. Briefly, 8–12-week-old *Prkcd*^*+/+*^ or *Prkcd*^−/−^ female mice were orthotopically implanted with E0771 cells (2.5 × 10^5^) as above. Mice from each genotype were randomized then treated intraperitoneally with anti-Ly6C or rat IgG2a (100μg/mouse) on day 0 followed by treatment with clodronate liposomes or control liposomes (200μl/mouse) according to manufacturer’s protocol on day 1. Anti-Ly6C or rat IgG2a treatments were given on days 0, 4 and 9, while clodronate or control liposomes were given on days 1, 5 and 10. Tumor volume was monitored until endpoint at day 14.

### *In vivo* macrophage co-injection studies

Primary bone marrow-derived macrophages (BMDM) from *Prkcd*^*+/+*^ or *Prkcd*^−/−^ female mice were polarized with IL-4 (20ng/mL) to M2-like phenotype for 24 hours and collected into a single cell suspension as previously described^43^. Purified cells were mixed 1:1 with LLC cells and 10^6^ total cells were injected subcutaneously into the right flank of naive eight-week-old C57BL/6J female hosts. LLC cells alone (10^6^) were used as a control. Tumor volume was measured every two days until endpoint.

### Isolation of single cells from mouse tumors

Excised tumors (~300 mg) were minced using scissors in RPMI media containing enzyme cocktail mix from Miltenyi Biotec mouse tumor dissociation kit (Miltenyi Biotec, Auburn, CA). Tumor pieces were further digested as per manufacturer’s instructions and digested tissue was filtered through 70μm strainer to obtain a single cell suspension. Spleen single cell suspensions were obtained by grinding spleens against a 70μm filter using a syringe plunger. Final single cell suspensions were obtained following red blood cell (RBC) lysis (Millipore Sigma, St. Louis, MO).

### Flow cytometry analysis

Flow cytometry was performed as described in our previous study^15^. Briefly, single cell viability was determined by using Ghost dye (Tonbo Biosciences Inc) followed by FcR-blocking (Tonbo Biosciences Inc). Antibodies were titrated, and the separation index was calculated using FlowJo v. 10 software (Treestar, Woodburn, OR). Cells were stained with fluorescently labeled antibodies as previously described^15^, and fixed with Foxp3/Transcription Factor Staining Buffer (Tonbo Biosciences). Stained cells were analyzed using Bio-Rad ZE5 flow cytometer in the UTHSC Flow Cytometry and Cell Sorting Core. A minimum number of 100 events were considered for analysis. Fluorescence minus one (FMO) stained cells and single color Ultracomp Beads (Invitrogen, Carlsbad CA) were used as negative and positive controls, respectively.

For *in vivo* intracellular staining, tumor single cell suspensions were stimulated with Cell Activation Cocktail (Biolegend) for 4 hours to allow accumulation of intracellular cytokines according to manufacturer’s protocol. After staining with cell surface markers, single cells were fixed and permeabilized with Flow Cytometry Perm Buffer (Tonbo Biosciences) followed by staining with IFNγ and TNFα.

Data were analyzed using FlowJo v. 10 software. Flow cytometry t-distributed stochastic neighbor embedding (tSNE) plots were generated using the built-in plugin in FlowJo to project and cluster gated flow cytometry immune cell populations^15^ (gating scheme shown in Supplementary Figure 1). All antibodies and reagents are provided in Supplementary Table 1.

### Isolation and stimulation of bone-marrow-derived macrophages, dendritic cells, and immature myeloid cells

Bone marrow cells were isolated from the femurs and tibias of *Prkcd*^*+/+*^ or *Prkcd*^−/−^ age-matched females and were cultured in complete RPMI media (5 × 10^5^ million cells/mL) supplemented with 50μM β-mercaptoethanol, 10mM HEPES, 1mM MEM non-essential amino acids (all Thermo Fisher). Bone-marrow-derived macrophages (BMDMs) were obtained after 6 days of culture with M-CSF (50ng/mL). BMDMs were left unstimulated or further polarized to an M1-like phenotype (M1 BMDMs) with IFNγ (20ng/mL) and LPS (100ng/mL) or to an M2-like phenotype (M2 BMDMs) with IL-4 (20ng/mL) for 24 h^62^. Bone-marrow dendritic cells (DCs) were obtained after 7 days of culture with FLT3L (100ng/mL) then left unstimulated or stimulated (DCstim) with LPS (100ng/mL) and anti-mouse agonistic CD40 monoclonal antibody (5μg/mL) for 24 h. Immature myeloid cells (iMCs) were obtained after culture with GM-CSF (40ng/mL) and IL-6 (40ng/mL).

### Antigen presentation and cross-presentation experiments and ELISA

CD4^+^ and CD8^+^ T cells were isolated from the spleens of tumor-free OT-II and OT-I mice, respectively, using magnetic-activated cell sorting (MACS, Miltenyi Biotec) according to manufacturer’s protocols and labeled with the proliferation dye CellTrace Violet (CTV). Purity was >90% for all populations as verified by flow cytometry analysis. BMDMs and DCs were pulsed with ovalbumin (OVA) (10μg/mL) for 24 h before co-culture with CD4^+^ (OT-II) and CD8^+^ (OT-I) T cells (105 cells) in a 96-well plate at a 1:2 antigen-presenting cell-T cell ratio for 72 h. Negative controls consisted of T cells cultured alone. T cell proliferation was assessed by CTV dilution within gated CD4^+^ and CD8^+^ T cells, respectively, and IFNγ levels were assessed by ELISA (R&D Systems, Minneapolis, MN) in the co-culture supernatants according to manufacturer’s protocol.

### RNA sequencing

*Prkcd*^*−/−*^ or *Prkcd*^*+/+*^ freshly isolated mouse BMDMs, M1 BMDMs, DCs, Dcstim and iMCs (n = 3 biological replicates each), as well as E0771 tumors (n = 5-6 biological replicates), were removed from dishes, and total RNA was collected using the the Rneasy Mini Kit (Qiagen) according to manufacturer’s instructions. The integrity of RNA was assessed using Agilent Bioanalyzer and samples with RIN > 5.0 were used. mRNA-seq libraries for the Illumina platform were generated and sequenced at GENEWIZ using the Illumina HiSeq 2×150bp configuration following manufacturer’s protocol.

### RNAseq analysis

Fastq files from Illumina HiSeq that passed quality control processing using FastQC^63^ were first aligned to the mouse transcriptome (mm10/GRCm38.p4 genome build with Ensembl v86 gene annotation) using STAR^64^ and then sorted with SAMtools^65^. Salmon^66^ was then used for transcript quantification and gene level counts were used for data analysis in R version 4.1.2^67^. Read counts were loaded from salmon quant files using tximport^68^, and differential gene expression analysis between *Prkcd^−/−^ and Prkcd^+/+^* groups was performed using DESeq2^69^. An adjusted p-value < 0.1 was used to determine significantly differentially expressed genes (DEGs) from each sample group described in the previous section. Read counts were normalized for downstream analyses and visualization using the variance stabilizing transformation (VST) from DESeq2. Heatmaps representing VST normalized and scaled gene expression values were generated with the ComplexHeatmap package^70^ where rows and/or columns were clustered via the “pearson” distance method. Significantly upregulated genes in *Prkcd*^*−/−*^ tumors, M1 BMDM, DCstim and iMCs were used as the input for the Gene Ontology (GO) Enrichment Analysis tool^71,72^. We performed a Bonferroni adjustment of gene set p-values for the number of gene sets tested in the GO software using Fisher’s Exact Test and biological processes were ranked by fold enrichment.

### Gene set enrichment analysis (GSEA)

For identification of enriched gene signatures, we used the GSEA software^73^. GSEA analysis was performed by using VST normalized gene expression data obtained from E0771 tumors, M1 BMDM, DCstim and iMCs (n = 5-6 for tumor and n = 3 biological replicates for other cell types). We used 1000 gene set permutations to test for significance at a false discovery rate (FDR) threshold of 0.25. The MSigDB hallmark gene sets (H collection)^74^ were used to determine enriched pathways in *Prkcd*^*−/−*^ and *Prkcd*^*+/+*^ groups. The top 10 ranked enriched genes by enrichment score in *Prkcd*^*−/−*^ groups are shown in a heatmap next to the corresponding GSEA enrichment plot (Figure 4 and supplementary figure 6).

### Single cell RNAseq analysis

PRKCD expression was analyzed in different immune cell populations within healthy or tumor human and mouse tissues using the online tool ‘Single Cell Portal’ from the Broad Institute https://singlecell.broadinstitute.org/single_cell). We used the following publicly available scRNAseq datasets: human triple negative breast cancer tumors (Wu et al., *EMBO J* 2020)^33^, human melanoma tumors (Jerby-Arnon et al., *Cell* 2018)^34^, human renal cell carcinoma tumors (Bi et al., *Cancer Cell* 2021)^35^, human colon cancer tumors (Pelka et al., *Cell* 2021)^36^, human glioblastoma tumors (Neftel et al., *Cell* 2019)^37^, human peripheral blood mononuclear cells (Broad/Boston and MtSinai/NYC) and mouse CD45^+^ splenocytes (Immgen labs).

### Statistical methods

Sample size for tumor studies were based on the effects observed in pilot studies and power calculations based on tumor growth studies. Power calculations were performed to ensure that the null hypothesis would be correctly rejected with > 80% power at 0.05 significance. For *in vivo* depletion and anti-PD-1 studies, mice were randomly assigned to experimental groups. Statistical differences between experimental groups were determined by unpaired Student’s t-tests for comparisons between two groups and one-way or two-way ANOVA with Tukey correction for multiple comparisons or two-way ANOVA with repeated measures to model longitudinal tumor growth between groups. Log-rank (Mantel-Cox) test was used to determine statistical significance for survival of mice. Statistical analysis was performed using the software within GraphPad Prism (GraphPad Software, Inc., La Jolla, CA). All data are shown as mean ± standard error of the mean (SEM). P values less than 0.05 were considered statistically significant.

## Funding

National Institutes of Health grant NCI R01CA253329 (LM)

National Institutes of Health grant NCI R37CA226969 (DNH, LM)

National Institutes of Health grant NCI UG1CA233333 (DNH)

National Institutes of Health grant NCI CA264021 (DNH)

The Mary Kay Foundation (LM)

V Foundation (DNH, LM)

ASPIRE Award from The Mark Foundation for Cancer Research (PGT)

National Institutes of Health NCI F31 CA268871 (MC)

Transdisciplinary Research on Energetics and Cancer R25CA203650 (LMS)

American Association for Cancer Research Triple Negative Breast Cancer Foundation Research Fellowship (LMS)

National Institutes of Health NCI F32 CA250192 (LMS)

The Obesity Society/Susan G. Komen Cancer Challenge award 2018 (LMS)

National Institutes of Health grant RO1 DK18061-01A1 (QW)

## Author contributions

Conceptualization: MC, PGT, DNH, LM

Formal analysis: MC, JH, MSB, LMS, TJH, LM

Investigation: MC, JH, LMS, JRY, MSB, UT, SJC, TJH

Methodology: MC, LMS, MSB, JH, SJC, QW, PGT, LM

Project administration: MC, LM

Resources: LM

Supervision: PGT, DNH, LM

Writing – original draft: MC, LM

Writing – review & editing: All authors

## Patient consent for publication

Not required

## Data availability statement

Data are available on reasonable request. All data relevant to the study are included in the article or uploaded as online supplementary information.

## Acknowledgments

We thank Dr. Hongbo Chi from the Department of Immunology at St. Jude Children’s Research Hospital in Memphis, TN for providing us with the spleens from OT-I and OT-II mouse spleens. We also thank Dr. Hasan Korkaya from the Georgia Cancer Center in Augusta, GA for proving us with the E0771-luc cell line. We thank Dr. James Carson from UTHSC Department of Medicine for providing us with LLC cells. We thank Drs. Zheng Dong and QingQing Wei from Augusta University in Augusta GA and Dr. Robert Messing from The University of Texas at Austin, Austin TX, for providing us with PKC delta knockout mice. Last, we thank the UTHSC Center for Cancer Research and the Flow Cytometry and Cell Sorting (FCCS) core at the University of Tennessee Health Science Center for their support.

---

**Supplementary Figure 1: Gating scheme for flow cytometry analysis**. Gating strategy for flow cytometric analyses of all myeloid cells and lymphocytes.

**Supplementary Figure 2: Antitumor immune pathways are enriched in *Prkcd*^*−/−*^ tumors**. **(A)** Heatmap of median-centered mRNA expression of differentially expressed genes (DEGs) in tumors from *Prkcd*^+/+^ and *Prkcd*^−/−^ mice. Adjusted p value less than 0.1. **(B-E)** GSEA plots for the GO gene sets of **(B)** T cell activation, **(C)** antigen processing and presentation, **(D)** innate immune response, and **(E)** inflammatory response in *Prkcd*^+/+^ and *Prkcd*^−/−^ E0771 tumors. (n = 5-6 biological replicates).

**Supplementary Figure 3: PKCδ expression in mouse and human tissues**. tSNE plots of single cells showing major cell types, PRKCD expression in **(A)** human renal cell carcinoma tumors (Bi et al., Cancer Cell 2021)^35^, **(B)** human colon cancer tumors (Pelka et al., Cell 2021)^36^, **(C)** glioblastoma tumors (Neftel et al., Cell 2019)^37^, **(D)** human peripheral blood mononuclear cells (Broad/Boston and MtSinai/NYC) and **(E)** *Prkcd* in mouse CD45^+^ splenocytes (Immgen labs).

**Supplementary Figure 4: Generation and activation of mononuclear phagocytes *in vitro*.** Experimental outline for **(A)** *in vitro* generation of different mononuclear phagocyte cell types and **(B)** activation of mononuclear phagocytes. GSEA of hallmark gene sets showing the most significantly enriched gene sets in **(C)** M1 BMDM compared to BMDM and **(D)** DCstim compared to DC are presented. Nominal p value <0.05.

**Supplementary Figure 5: Response to Type I IFN in *Prkcd*^*−/−*^ mononuclear phagocytes**. Heatmap of median-centered mRNA expression of genes from hallmark response to type I interferon in tumors from *Prkcd*^+/+^ and *Prkcd*^−/−^ mice (n = 5-6 biological replicates).

**Supplementary Figure 6: PKCδ promotes protumor and anti-inflammatory pathways in mononuclear phagocytes**. GSEA of hallmark gene sets showing the most significantly enriched gene sets in *Prkcd*^+/+^ **(A)** E0771 tumors, **(B)** iMCs, **(C)** M1 BMDMs and **(D)** DCstim. **(E)** Venn diagrams of commonly enriched hallmark pathways between *Prkcd*^+/+^ tumors, M1 BMDMs, DCstim and iMCs (n = 3 biological replicates for all mononuclear phagocytes and n = 5-6 for E0771 tumors). Nominal p value < 0.05.

**Supplemental Table 1.**
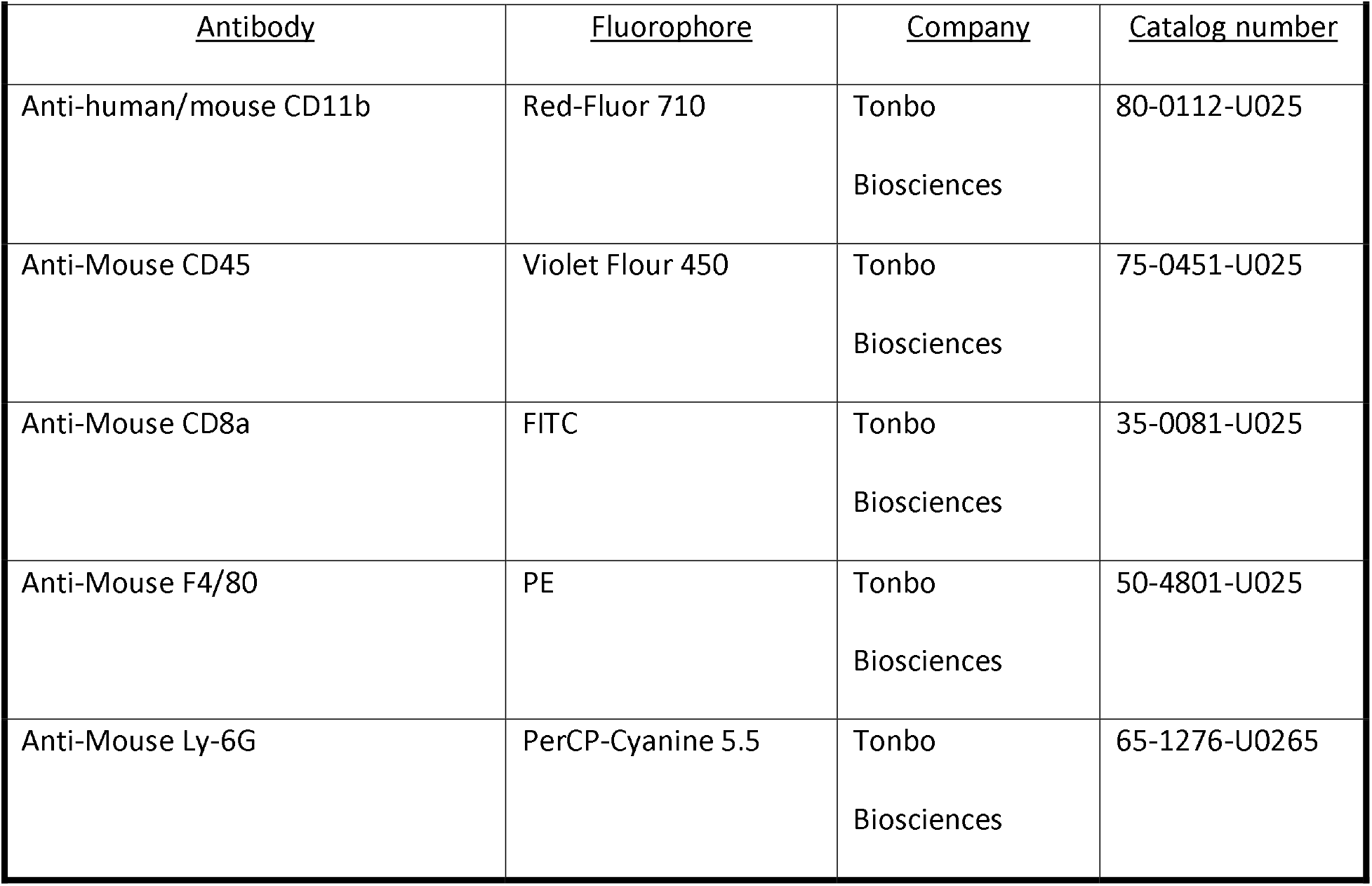
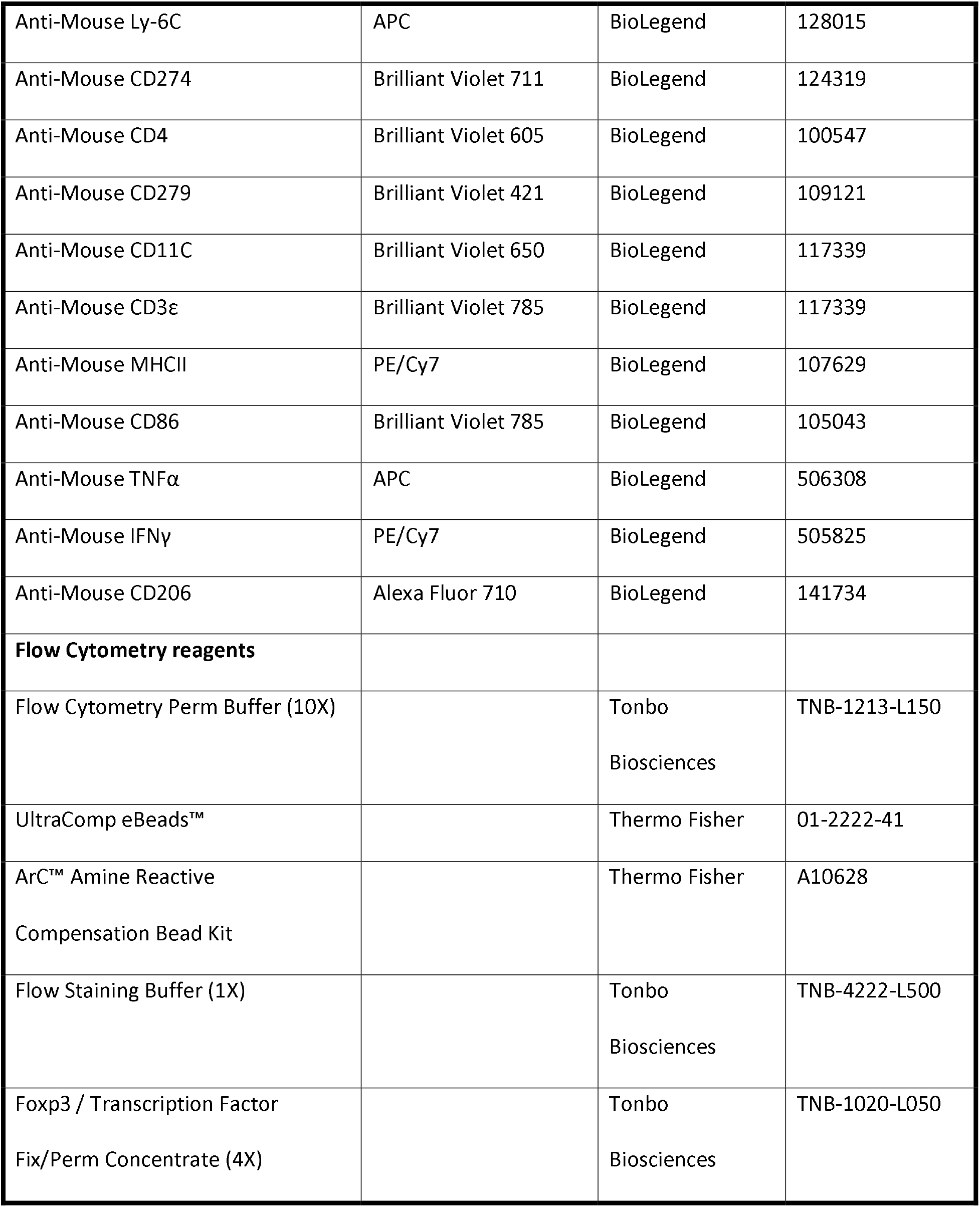
Flow cytometry antibodies and reagents.

