## Supplementary materials for "Protein Kinase C Delta Regulates Mononuclear Phagocytes and Hinders Response to Immunotherapy in Cancer"

**Supplementary Figure 1: Gating scheme for flow cytometry analysis**. Gating strategy for flow cytometric analyses of all myeloid cells and lymphocytes.

**Supplementary Figure 2: Antitumor immune pathways are enriched in *Prkcd^-/-^* tumors**. **(A)** Heatmap of median-centered mRNA expression of differentially expressed genes (DEGs) in tumors from *Prkcd*^+/+^ and *Prkcd*^-/-^ mice. Adjusted p value less than 0.1. **(B-E)** GSEA plots for the GO gene sets of **(B)** T cell activation, **(C)** antigen processing and presentation, **(D)** innate immune response, and **(E)** inflammatory response in *Prkcd*^+/+^ and *Prkcd*^-/-^ E0771 tumors. (n = 5-6 biological replicates).

**Supplementary Figure 3: PKCδ expression in mouse and human tissues**. tSNE plots of single cells showing major cell types, PRKCD expression in **(A)** human renal cell carcinoma tumors (Bi et al., Cancer Cell 2021)^35^, **(B)** human colon cancer tumors (Pelka et al., Cell 2021)^36^, **(C)** glioblastoma tumors (Neftel et al., Cell 2019)^37^, **(D)** human peripheral blood mononuclear cells (Broad/Boston and MtSinai/NYC) and **(E)** *Prkcd* in mouse CD45^+^ splenocytes (Immgen labs).

**Supplementary Figure 4: Generation and activation of mononuclear phagocytes *in vitro*.** Experimental outline for **(A)** *in vitro* generation of different mononuclear phagocyte cell types and **(B)** activation of mononuclear phagocytes. GSEA of hallmark gene sets showing the most significantly enriched gene sets in **(C)** M1 BMDM compared to BMDM and **(D)** DCstim compared to DC are presented. Nominal p value <0.05.

**Supplementary Figure 5: Response to Type I IFN in *Prkcd^-/-^* mononuclear phagocytes**. Heatmap of median-centered mRNA expression of genes from hallmark response to type I interferon in tumors from *Prkcd*^+/+^ and *Prkcd*^-/-^ mice (n = 5-6 biological replicates).

**Supplementary Figure 6: PKCδ promotes protumor and anti-inflammatory pathways in mononuclear phagocytes**. GSEA of hallmark gene sets showing the most significantly enriched gene sets in *Prkcd*^+/+^ **(A)** E0771 tumors, **(B)** iMCs, **(C)** M1 BMDMs and **(D)** DCstim. **(E)** Venn diagrams of commonly enriched hallmark pathways between *Prkcd*^+/+^ tumors, M1 BMDMs, DCstim and iMCs (n = 3 biological replicates for all mononuclear phagocytes and n = 5-6 for E0771 tumors). Nominal p value < 0.05.

| **Supplemental Table 1**. **Flow cytometry antibodies and reagents** | | | |
| --- | --- | --- | --- |
| Antibody | Fluorophore | Company | Catalog number |
| Anti-human/mouse CD11b | Red-Fluor 710 | Tonbo Biosciences | 80-0112-U025 |
| Anti-Mouse CD45 | Violet Flour 450 | Tonbo Biosciences | 75-0451-U025 |
| Anti-Mouse CD8a | FITC | Tonbo Biosciences | 35-0081-U025 |
| Anti-Mouse F4/80 | PE | Tonbo Biosciences | 50-4801-U025 |
| Anti-Mouse Ly-6G | PerCP-Cyanine 5.5 | Tonbo Biosciences | 65-1276-U0265 |
| Anti-Mouse Ly-6C | APC | BioLegend | 128015 |
| Anti-Mouse CD274 | Brilliant Violet 711 | BioLegend | 124319 |
| Anti-Mouse CD4 | Brilliant Violet 605 | BioLegend | 100547 |
| Anti-Mouse CD279 | Brilliant Violet 421 | BioLegend | 109121 |
| Anti-Mouse CD11C | Brilliant Violet 650 | BioLegend | 117339 |
| Anti-Mouse CD3ε | Brilliant Violet 785 | BioLegend | 117339 |
| Anti-Mouse MHCII | PE/Cy7 | BioLegend | 107629 |
| Anti-Mouse CD86 | Brilliant Violet 785 | BioLegend | 105043 |
| Anti-Mouse TNFα | APC | BioLegend | 506308 |
| Anti-Mouse IFNγ | PE/Cy7 | BioLegend | 505825 |
| Anti-Mouse CD206 | Alexa Fluor 710 | BioLegend | 141734 |
| **Flow Cytometry reagents** |  |  |  |
| Flow Cytometry Perm Buffer (10X) |  | Tonbo Biosciences | TNB-1213-L150 |
| UltraComp eBeads™ |  | Thermo Fisher | 01-2222-41 |
| ArC™ Amine Reactive Compensation Bead Kit |  | Thermo Fisher | A10628 |
| Flow Staining Buffer (1X) |  | Tonbo Biosciences | TNB-4222-L500 |
| Foxp3 / Transcription Factor Fix/Perm Concentrate (4X) |  | Tonbo Biosciences | TNB-1020-L050 |
