## Supplementary figures for "Protein Kinase C Delta Regulates Mononuclear Phagocytes and Hinders Response to Immunotherapy in Cancer"

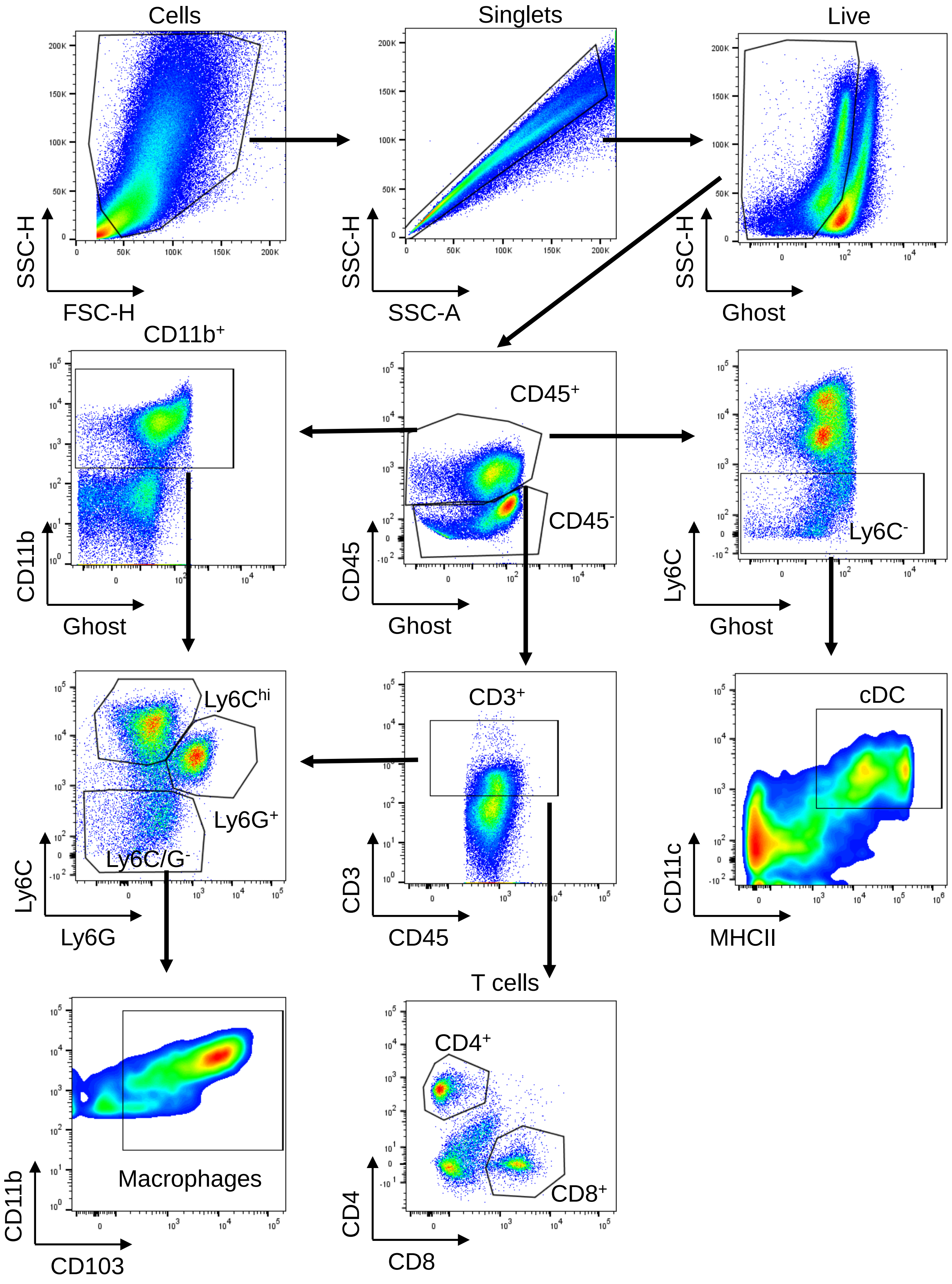


**Supplementary Fig 1**

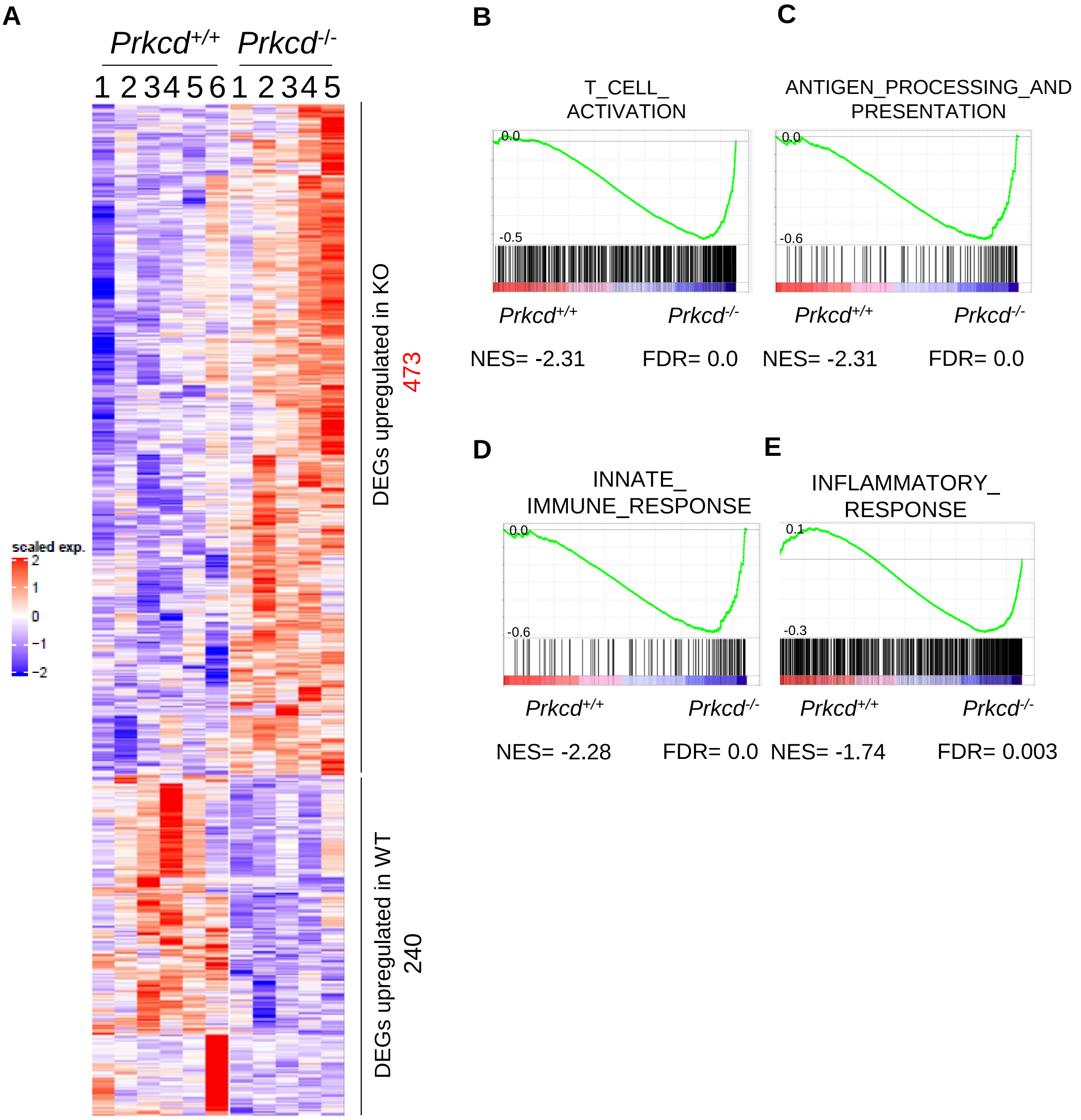


**Supplementary Fig 2**

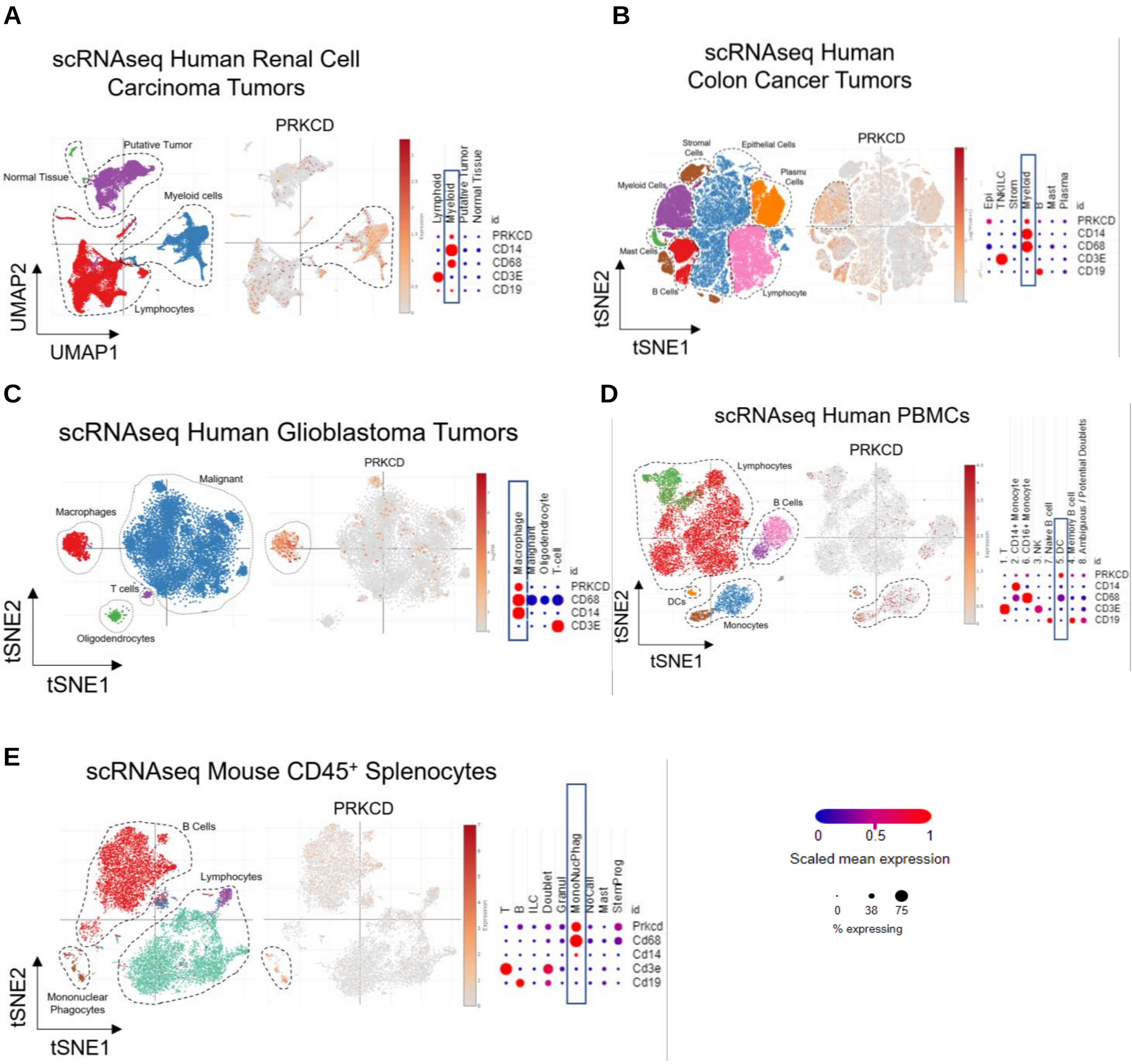


### Supplementary Fig 3

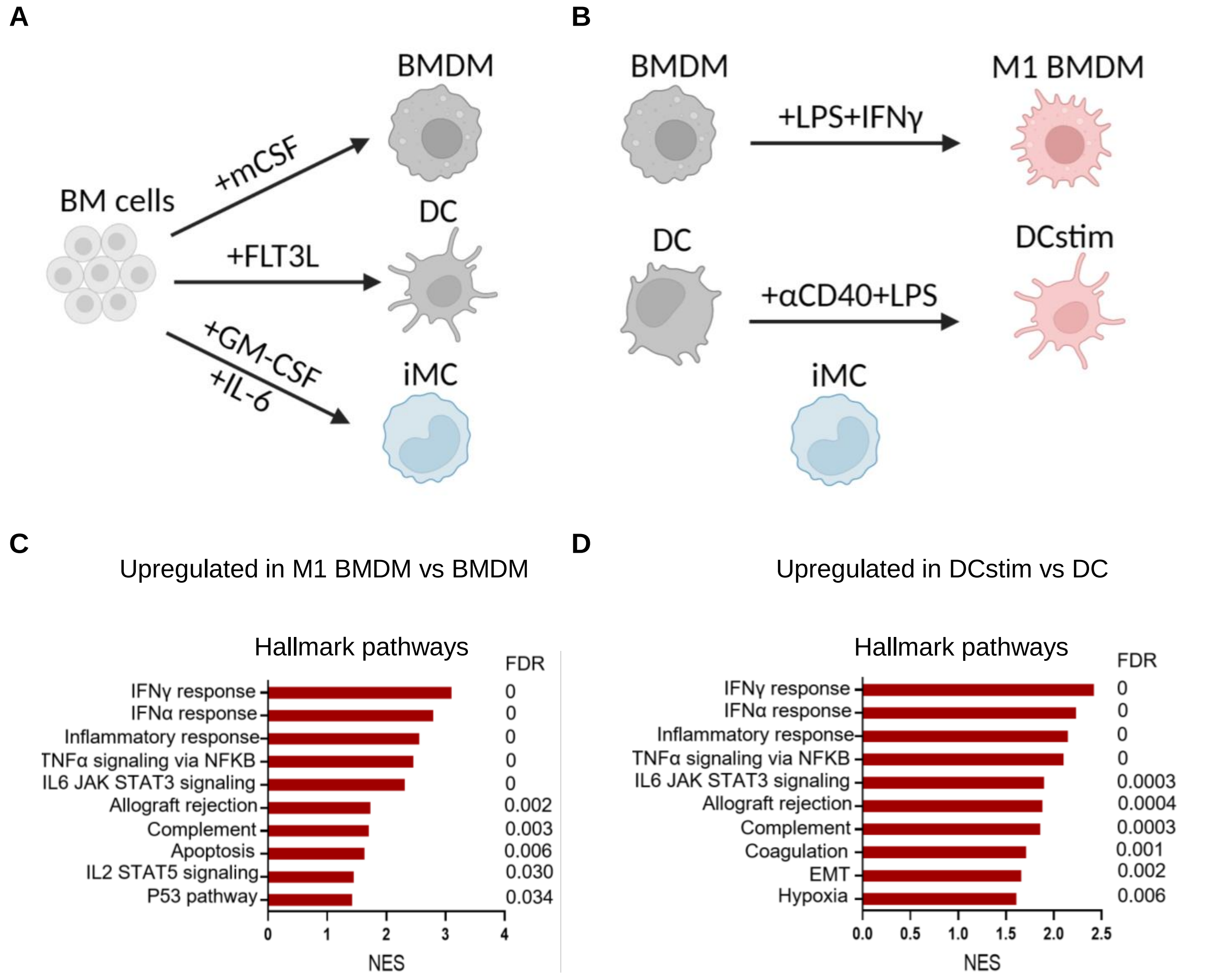


### Supplementary Fig 4

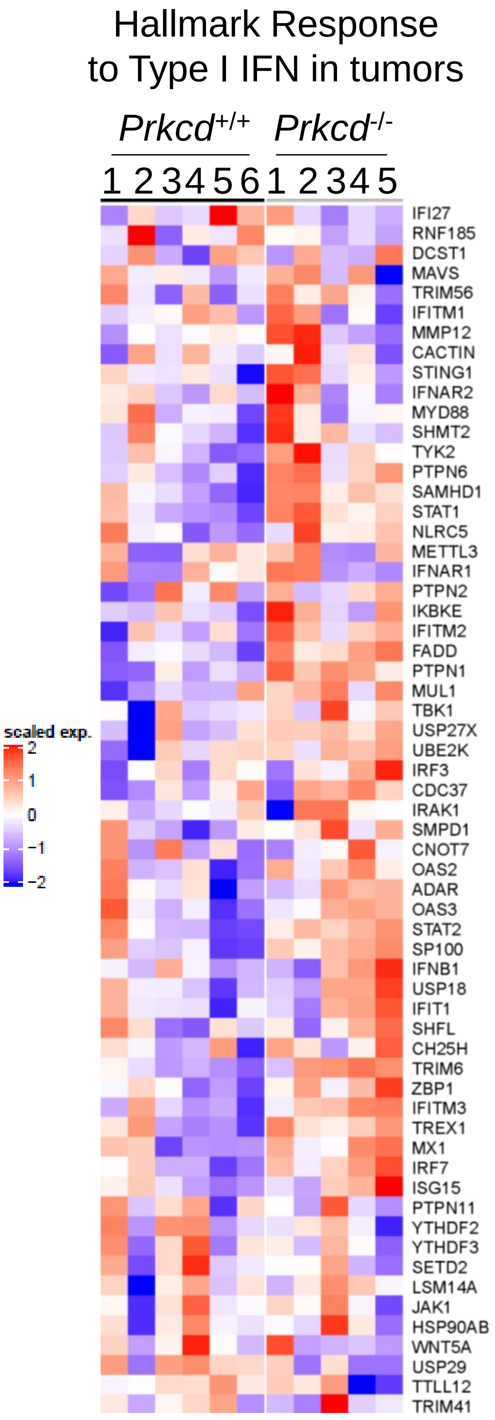


**Supplementary Fig 5**

**Hallmark pathways enriched in *Prkcd*$^{+/+}$**
Nominal p value < 0.05

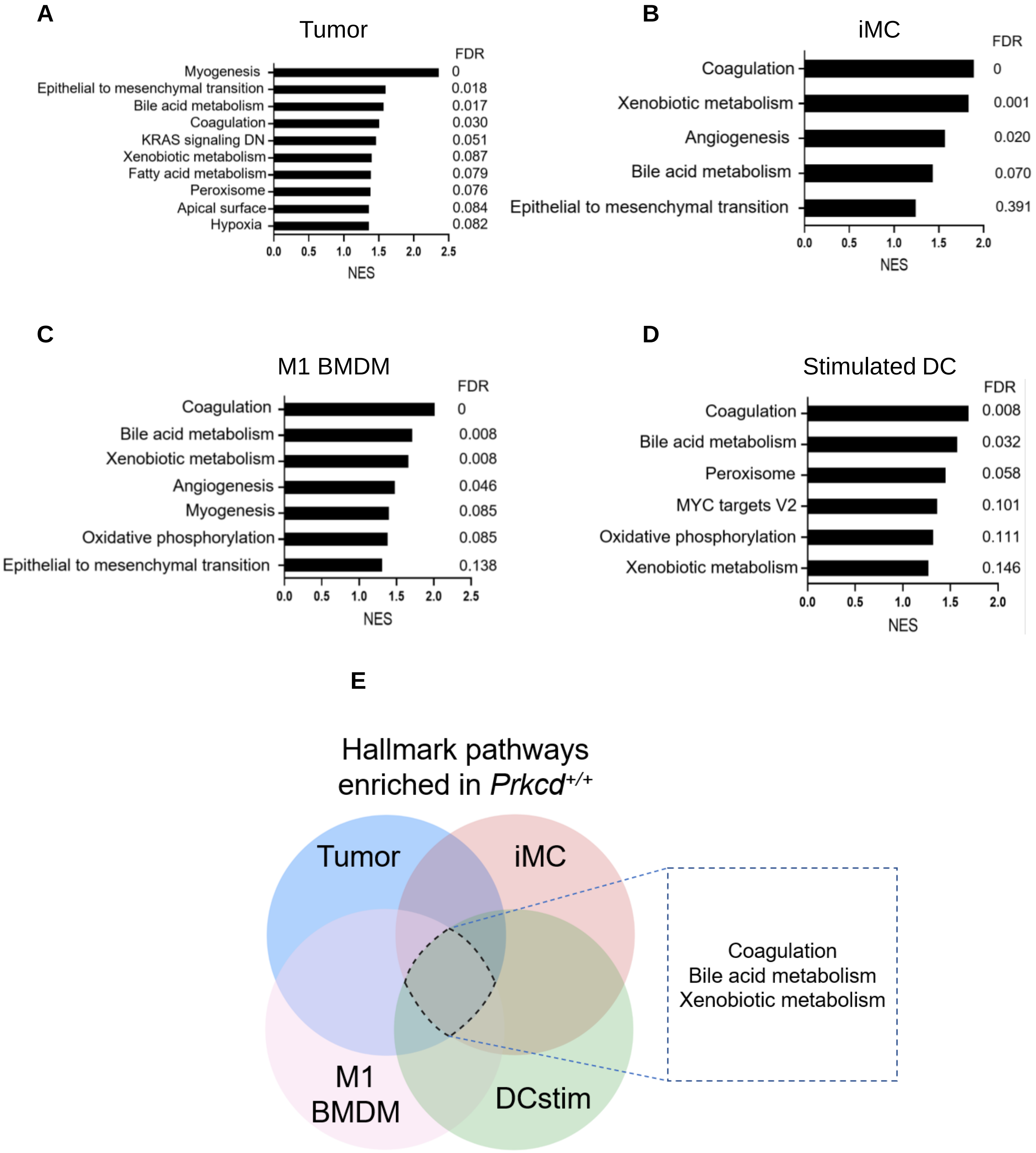


**Supplementary Fig 6**
